# Forest cover is more important than habitat integrity or landscape configuration in determining habitat use by mammals in a human-modified landscape in Colombia

**DOI:** 10.1101/2023.02.27.530271

**Authors:** Lain E. Pardo, Bibiana Gómez-Valencia, Nicolas J. Deere, Yenifer Herrera Varón, Carolina Soto, Elkin A. Noguera-Urbano, Lina M. Sánchez-Clavijo, Luis Romero, Angélica Díaz-Pulido, José Manuel Ochoa Quintero

## Abstract

Human activities shape the structure of landscapes in different ways and hence modify animal communities depending on the type and intensity of these activities. The Montes de Maria subregion of Colombia has experienced a heavy transformation of most areas despite covering one of the last remnants of dry forest, a critically endangered ecosystem. However, the effects of this transformation have been little explored. Here, we used a multispecies occupancy model (MSOM) to understand the relative influence of three components of land-use change – deforestation (remaining habitat amount), degradation of habitat quality (habitat quality) and fragmentation (landscape configuration) on mammalian habitat use across a mosaic of tropical dry forest in Colombia. Our data suggest that the percentage of forest cover was substantially important for herbivores, and consistently showed a moderate effect on the entire community and some individual species. High variability in species-specific responses to the examined variables hindered broad taxonomic generalizations, nevertheless, we detected a moderate positive effect of forest cover in both diet specialists and generalists species, as well as in species with small home ranges. Although omnivores responses, tended to use less complex landscapes (mosaics of land uses), there was high uncertainty in this response. The lack of substantial effects on most species, and the absence of threatened species across this anthropogenic landscape, suggests that the current community is composed of species tolerant to habitat modifications, but not only diet generalist species. This is most likely the result of a long filtering process caused by land use transformation and hunting which could have caused non-sensitive species to distribute relatively homogenously across this landscape. Our results suggest that conservation strategies in the study area should focus on conserving and expanding as much forest as possible rather than only improving the quality of already existing forest patches.

## 1. Introduction

Habitat loss and fragmentation (splitting of habitats, sensu Fahrig, 2003) are well-known consequences of human activities that strongly affect biodiversity and ecosystem function in different ways (Crooks et al., 2017; Dirzo et al., 2014). Human activities negatively affect most vertebrate species by limiting their movements and creating ecological traps with an elevated risk of mortality, particularly for rare and specialized species (Battin, 2004) and reducing species richness and abundance (Deere et al., 2020; Fahrig, 2003). Although mammals play important roles as herbivores, predators, ecosystem engineers, and keystone species (Lacher et al., 2019), at the same time, they are among the most threatened groups by human-driven processes (Ceballos et al., 2017). However, their responses can vary between species with some species being more tolerant than others to anthropogenic environments such as agroecosystems (Pardo et al., 2018a) or urban environments (Hansen et al., 2020). Fragmentation processes affect not only taxonomic level parameters such as species richness but also other components such as the trophic structure of a community, functional traits, distribution of populations, or species interactions, among other processes (Magioli et al., 2016; Morris, 2010). Evidence suggests, for example, that current anthropogenic pressures tend to homogenize vertebrate communities by a filtering mechanism which can result in communities mainly consisting of disturbance-tolerant generalist species with wide niche breaths or reduced functional traits and networks (McKinney and Lockwood, 1999; Püttker et al., 2015). For example, Lomolino and Perault, (2007) found that the body mass of small mammals tends to be lower in disturbed/fragmented habitats than in undisturbed habitats Similarly, infrastructure seems to promote smaller species with less carnivorous diets (Li et al., 2021; Suraci et al., 2021).

Recently, important discussions have taken place regarding which factors are more important in determining biodiversity and community structure of animals. In particular, whether quality or quantity is more important for species distribution or survival. Regolin et al. (2021), for example, suggest that quality is more important than forest cover for the habitat use of some neotropical mammals. On the other hand, (Fahrig, 2013) suggests in her “habitat amount hypothesis” that the quantity of forest cover (not the number of patches or individual size) is more important regardless of the configuration of the system (a fragmentation surrogate), while Zungu et al (2020) found that both habitat amount and its spatial configuration were important for the occurrence of mammals. Since all responses can vary locally and depending on the intensity of habitat modification, it is important to assess these relationships in different contexts or biomes. In human-dominated landscapes, species’ responses to abiotic and biotic factors determine which species persist in the landscape. Therefore, understanding the relative contribution of different components of land use change or habitat modification on mammal populations is crucial to visualize conservation strategies. For example, knowing how and which animal species persist in disturbed landscapes is important not only for protecting rare populations or controlling potential problematic species but also because of their functional roles in the ecosystem (Newbold et al., 2018).

Biodiversity assessments usually focus on either species richness or abundance. While these quantities can provide important indications of the system’s state, some challenges are associated when using these metrics. For example, although abundance is the most relevant parameter for monitoring biodiversity, it is very difficult to measure, particularly for species that do not possess unique markings. On the other hand, species richness alone is limited as an indicator of biodiversity change because taxonomic identities and species-specific responses are not accounted for (Hillebrand et al., 2017; Pardo et al., 2018a). Furthermore, diversity includes many other important aspects, such as trophic structure, functional traits, phylogenetic relationships of species, and the distribution of individuals within a community across the landscape (Noss, 1990). Functional diversity, in particular, is a recent popular metric that allows us to understand how the biological traits of species, such as trophic level or guild, and phylogenetic relationships among others, influence the functioning of ecosystems (Tilman, 2001). For example, Suraci et al. (2021) found that life history traits were important in a North American landscape, suggesting that human presence and infrastructure promote smaller species with less carnivorous diets. Although there is increasing information about trait types for mammals (Jones et al., 2009; Wilman et al., 2014), the relationship between these traits and responses to environmental variables is still little explored, especially in anthropogenic landscapes with growing human pressures.

In this paper, we aim to understand the relative contribution of different components of habitat modification on mammal communities across the Montes de Maria subregion in Colombia. Specifically, we were interested in understanding how the entire community, three functional traits (trophic guild, diet breadth, and home range), and individual species respond to the degradation of habitat quality, remaining habitat amount (extent of forest), and fragmentation. For this, we used four surrogates: 1) forest integrity as an index of anthropogenic alteration of habitat quality; 2) forest cover as a measure of remaining habitat availability which at the same time provides an indication of deforestation processes; 3) conditional entropy and 4) core area metrics as measures of fragmentation processes (i.e. complexity of the landscape, and landscape configuration, respectively). Based on current evidence, we hypothesized that most species occurring in this disturbed region are not drastically affected by fragmentation per se due to long-term selective pressure by humans (filtering), which would have resulted in only disturbance-tolerant species occupying the region. However, the habitat use by species would be mostly dependent on the percentage of the available forest as suggested by the habitat amount hypothesis (Fahrig, 2013). As such, we predicted that the percentage of forest cover will have stronger effects than metrics of configuration and integrity. However, these responses would vary among species and functional groups, with more specialized guilds and species with small home ranges being more dependent on the integrity of the forest.

Our appraisal focuses on the Colombian-Caribbean region. This region faces great conservation and social challenges (Negret et al., 2019) where deforestation, agriculture, livestock, illegal activities, and unauthorized burning have increased considerably (Etter et al., 2008).These processes have had an important effect on the relationships between species and the functioning of the original ecosystems. In fact, Colombia has lost around 92% of the original dry forest (lowland forest with high strong rainfall pattern), and only fragmented areas remain (Garcia et al., 2014). The Caribbean region, in particular, has lost about 45% of the original dry forest. As such, the “Montes de Maria” subregion, where this study took place, is one of the most important for biodiversity in the Caribbean and Colombia since it is one of the last few places where some patches of Tropical Dry Forest still present, despite the heavy transformation of most areas and social conflicts (Garcia et al., 2014). Due to its social and ecological importance, current conservation efforts coincide in that one of the largest gaps of information for decision-making is understanding the relationship between fauna and its habitats and its repercussions on ecosystem services (Norden et al., 2020). Understanding which landscape factors are more important than others helps to inform conservation practice. For example, whether efforts in restoration should focus on improving quality or increasing the amount of habitats (forests) to promote conservation.

## 2. Materials and Methods

### 2.1. Study Area

The study area corresponds to the Montes de María subregion in the Colombian Caribbean, encompassing the areas between the municipalities of San Juan Nepomuceno (Department of Bolivar) and Colosó in the Department of Sucre (Geographic coverage: minimum latitude and minimum longitude [9.5, -75.37], maximum latitude and maximum longitude [10.04, -75.05]). The area is a heterogeneous anthropogenic landscape, comprised of tropical dry forest fragments of different ages or successional stages, shrubby vegetation, permanent and seasonal small holding agricultural crops (ñame, cacao, teak, oil palm, tobacco, other fruits) and deforested areas for ranching activity. Montes de María was one of the regions of Colombia where biodiversity has been significantly altered over the course of the last century, due to habitat loss, fragmentation and illegal wildlife trade.

### 2.2. Data Collection

We surveyed 89 sites across the Magdalena Medio subregion using a single camera trap per site (Bushnell 24MP Aggressor Low Glow HD) from July to September 2018. Before deploying the cameras, we divided the study area into ∼98 1200 ha hexagon-shaped polygons representing a gradient of forest cover (Fig. 1) using ArcGis (ESRI, 2011). Due to security concerns and accessibility constraints, we selected 12 hexagons for analysis. Therefore, our sampling unit is the camera site inside each of these hexagons. These units were distributed in the two sectors, the North covering approximately 13,203 ha (minimum convex polygon with 44 camera sites) and the South covering 6,434 ha (45). The north area is close to Los Colorados Fauna and Flora Sanctuary, while sites in the southern part are near a small regional protective forest (Serranía de Coraza Protective Forest Reserve). We implemented a systematic survey design whereby the first camera was placed randomly and subsequent cameras were separated by an inter-trap distance of 1 km. Fine-scale deployments were guided by evidence of animal activity (e.g. trails, footprints, faeces, among others) and, in some circumstances human-made trails. Cameras were left in the field for approximately 60 days (mean = 54 days per camera, range = 12-63) secured to trees at a height of 30-60 cm and no bait or lures were used during the deployment procedure. Cameras provided continuous 24-hour monitoring of the sampling location and were programmed to capture a combination images and videos with 2-seconds intervals and high sensor sensitivity.

**Figure 1.**
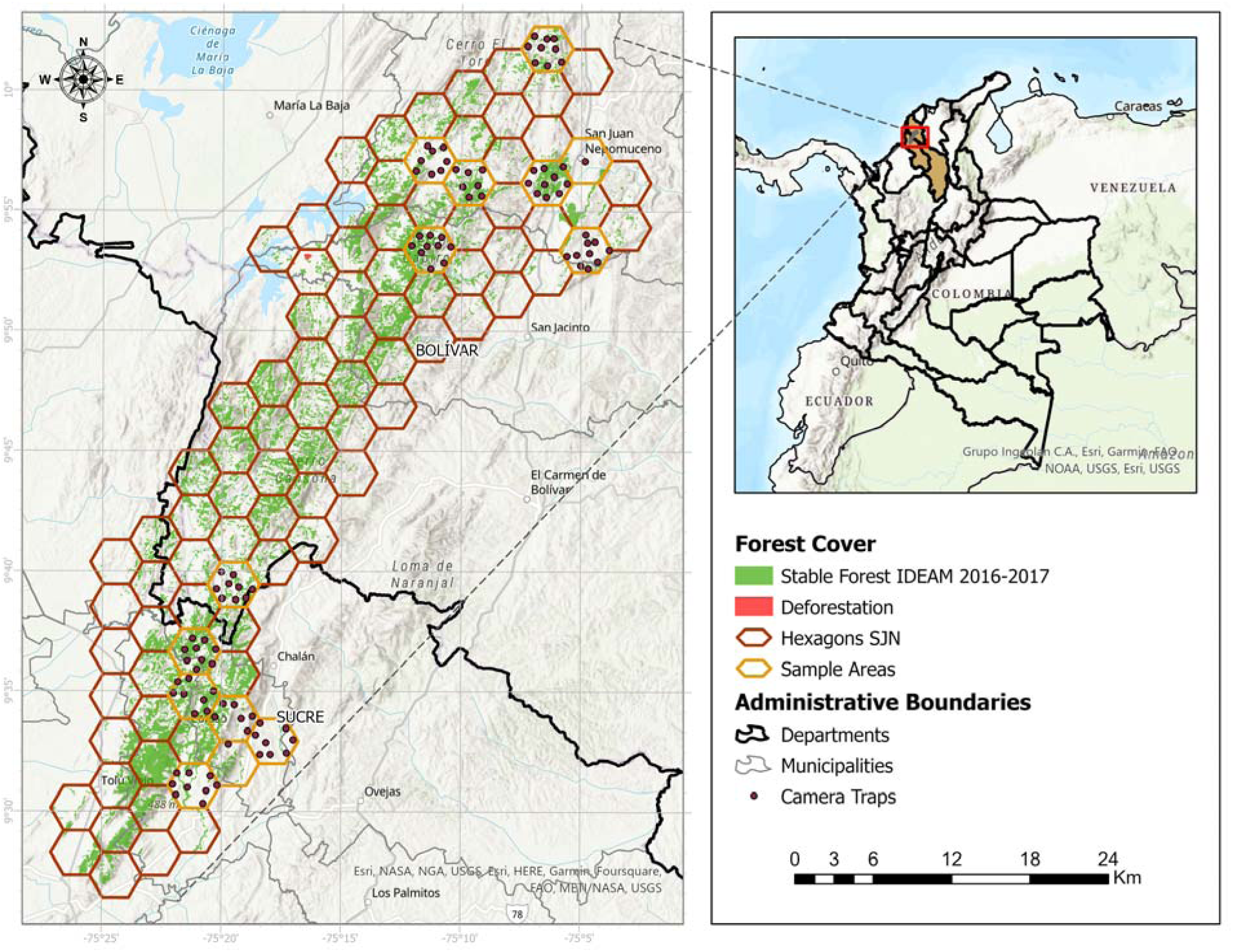
Study area and sampling design in the Montes de Maria subregion (Colombia). Hexagons show the preliminary division of the area where camera traps were deployed along a forest cover gradient.

### 2.3. Landscape Features

We used four covariates that we hypothesized could have an important effect on mammal habitat use: Forest Landscape Integrity Index (FLII), which shows the conservation status of the forest by measuring the degree of forest modification for the beginning of 2019 (Grantham et al., 2020), and hence, here we used it as an index of forest quality. Forest cover (FOREST) as a measure of habitat extent or availability, Conditional Entropy (ENTROPY), and the Coefficient of variation of core area (COREAREA), both as measures of landscape configuration (complexity and patch shape/area, respectively). FLII combines observed pressures, inferred pressures and loss of forest connectivity, and the resulting layer has a spatial resolution of 300 m, ranging from 0 to 10, and the NA zones are non-forest zones defined (see Grantham et al., 2020). The higher values of FLII represent forests with greater integrity (i.e. the degree to which its structure, composition and function have been modified by anthropogenic actions; sensu Parrish et al., 2003). Two components of this index (observed human pressures and inferred human pressures) represent the human footprint that influences the integrity of forests, such as current land use, population density, intervention time, distance to roads, and a forest-adjusted biomass index (the value used corresponded to the average of these values inside the buffer). Throughout, we extracted all covariates as average values across buffers of 500 m radius around each camera trap and interpreted these areas as landscapes (i.e. landscapes of 78 ha).

Forest cover is simply the percentage of forest cover within the buffer. Conditional Entropy represents the complexity of the configuration of a spatial pattern. If the value of the conditional entropy is low, the cells of one category have adjacent cells predominantly of one category. However, if the value is high, it means that cells from one category have adjacent cells from many categories (Nowosad and Stepinski, 2019). Core area describes patch area and shape simultaneously (more core area when the patch is large and the shape is rather compact, i.e. a square) (Hesselbarth et al., 2019). To calculate the above variables, we used the Corine Land cover shapefiles for Colombia (IDEAM et al., 2007), which are available at a 30 m resolution. Landscape variables were calculated and extracted with ArcGIS*®* and the software R (R Core Team, 2020) using the package Landscape metrics (Hesselbarth et al., 2019). We also considered six other landscape variables (Mean edge density of forest classes, Simpson’s Landscape Diversity Index, percentage of agriculture, mesh size, Simpson evenness, Shannon entropy). However, after examining for collinearity, using Spearmańs rank correlation coefficient (>70%), these variables were omitted from the analysis due to multicollinearity issues (these metrics were also calculated for the buffer of 500 m radius).

### 2.4. Traits Responses

We analyzed mammal responses to covariates at three taxonomic scales: community-level, species-level and functional/traits groups (trophic guild, dietary breadth, and home range). To determine the trophic guild of mammals and home range categories, we used the EltonTraits database (Wilman et al., 2014) to determine the percentage of food items consumed by each species (e.g, plant material, grass, vertebrates, etc.). Once these percentages were annotated, we classified the species into four groups following Oberosler et al. (2019) approach: (1) carnivore (>50% of diet based on vertebrates), (2) herbivorous (includes herbivores, browsers, granivores, and frugivores, with >50% plant material), (3) insectivorous (>50% invertebrates), (4) omnivorous (usually both plant and animal material). Wilman et al. (2014) define home range as the area (km^2^) within which everyday activities of individuals or groups are typically restricted, estimated by either direct observation, radio telemetry, trapping or unspecified methods over any duration of time (see Wilman et al 2014 for details) For taxa that could not be reliably identified to species-level, functional/traits characteristics were derived from the species’ closest relative or the genus (e.g, “*Mazama* sp” and “*Coendu* sp”).

To determine the dietary breadth, we reclassified the PanTHERIA database (Jones et al., 2009) into wider categories for ease of analysis. Original values indicated a range of dietary breadth from 1-7, so we redistributed this into only two categories: narrow diet (1-3), representing specialist species, and wide diet (4-7), representing dietary generalists. Dietary breadth represents the “number of dietary categories eaten by each species measured using any qualitative or quantitative dietary measure over any period of time, using any assessment method, for non-captive or non-provisioned populations; adult or age-unspecified individuals, male, female, or sex-unspecified individuals; primary, secondary, or extrapolated sources; all measures of central tendency; in all localities” (see Jones et al 2009). Dietary categories were defined as vertebrate, invertebrate, fruit, flowers/nectar/pollen, leaves/branches/bark, seeds, grass, and roots/tubers. For detail of this classification (see Appendix A.1).

### 2.5. Modeling Framework

We analyzed the variation in mammal species richness and occupancy (state variables) across the Montes de Maria region using Bayesian hierarchical multi-species occupancy models (R. Dorazio & Royle, 2005) with parameter-expanded data augmentation (see also Dorazio et al., 2006; Zipkin et al., 2010). This hierarchical framework explicitly accounts for the ecological process (occupancy) and the observational process (detection) which makes it a robust framework for reducing the bias in the inferences due to false-negative measurement errors. It also allows estimating occurrence probabilities at different taxonomic scales (i.e. community and individual species) for multiple species simultaneously, as species-specific parameters are treated as random effects, drawn from a common distribution, characterized by estimable hyperparameters that represent the overall response of the mammal community. This helps to improve inferences for rare species with low detections due to the borrowing of information by individuals across the community (Kéry and Royle, 2016). Previous to the modeling, we standardized covariates (subtracting the mean and dividing by the standard deviation).

The observed data (Y_ij_) corresponds to the number of sampling occasions out of a total of *k_j_* sampling occasions that species *i* was detected at camera station *j.* We grouped the detections per day into 10 sampling occasions. Only species detected at least on two sampling occasions were included (MacKenzie, 2006). We did not filter rare species from the community as we did not want to lose species from the different functional groups we were examining and therefore suggest caution in interpreting the individual species-specific occupancy associations of species with less than five detections or detected in less than 10% of the sites (i.e. Cougar– *Puma concolor*, Brown-eared woolly opossum– Caluromys lanatus, Brown four-eyed opossum– *Metachirus nudicaudatus*, Greater grison– *Galictis vittata*, Margay– *Leopardus wiedii*, Eastern cottontail–*Sylvilagus floridanus*, coendu–*Coendou* sp).

#### Model Formulation

We modelled the latent occupancy state, z_i,j_, of species *i* at site *j* (ψ _i,j_) as a Bernoulli process, z_i,j_ ∼ Bern(ψ_i,j_), where ψ_i,j_ denotes the probability of occurrence(ψ). We estimated the observation process using a second Bernoulli process, x_i,j,k_ ∼ Bern(p_i,j,k_* z_i,j_), where x_i,j,k_ represents the observed detection/non-detection data of species *i*, at site *j*, during sampling occasion *k*, and p_i,j,k_ is the probability of detecting the species conditional on its presence at the site (z_i,j_ = 1) (Zipkin et al., 2010).The model assumes that variation in the abundance of a species across sampling sites does not affect species detection probabilities p_i,j,k_ . We specified models of the form:

#### Occupancy model (ecological process)

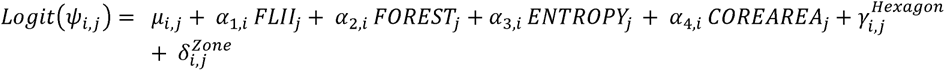

#### Observational model (Detection)

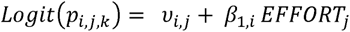

Occupancy and detection probabilities were modelled with species-specific intercepts (µ and v respectively) on the logit scale. We described occupancy (ψ _i,j_) as a function of habitat extent, quality and configuration (α_1-4_). Due to inherent spatial clustering in the survey design, we implemented random effects to account for potential lack of independence between cameras in hexagonal sampling units (*y_i,j_^Hexagon^*) and zones (δ*_i,j_^Zone^*). For the zonal random effects, we used a Half-Cauchy prior to account for the low number of factor levels (N=2). These two zones can represent different situations, with the north area being more conserved than the south area, and therefore could be used as a proxy for “conservation status”. We described detection probability as a function of survey effort (the number of days the camera was active; EFFORT; β_1_). We used the posterior draws to estimate the effect of the variables on functional groups (trophic guild, diet breadth, and home range, see Appendix B for details on the grouping).

All analyses were performed in R (version 4.3.0, R Core Team, 2020) using JAGS via the ‘jagsUI’ package (Kellner, 2021), applying three parallel MCMC chains (nc), with 150,000 iterations (ni), removing the first 50,000 as a period of burn-in (nb), posterior chains were thinned by 100 (nt). We constructed models using uninformative priors, including diffuse normal priors for intercept and slope parameters and wide uniform priors for variance parameters. We used the Gelman diagnostic plots to assess model convergence visually (Gelman et al., 1996). We further used two measures of model fit for the posterior predictive distribution using the lack of fit statistic (chat) and Bayesian P value (Bpv) with quantities close to 0.5 indicating adequate model fit, while 0.05 < P > 0.95 indicative of poor model fit. Throughout, we considered covariates to have a substantial effect if the 95% Bayesian credible interval (BCI) of the associated parameter did not overlap zero, and a moderate effect if 75% BCIs did not overlap zero.

#### Species Richness

We used data augmentation as described by Kéry & Royle, (2016) to estimate species richness of the entire community as a derived parameter. This process appends a number of hypothetical species with all zero encounter histories to the detection matrix to estimate the number of species that occupied the study landscape(s) but were undetected by the survey protocols. We estimate that there could be up to 46 terrestrial mammal species detectable by camera trapping in the study area, based on their distribution knowledge. Given that 21 species were represented in our detection matrix, we use M (“true” species) = 46 minus n (observed species) as our data augmentation parameter (25 potentially undetected species). Implementation of the model with a uniform prior is done by augmenting the data set with M-n all-zero encounter histories. Then the model for the augmented data set is a zero-inflated version of a model where the actual number of species in the community (N) is known (see also (Zipkin et al., 2010) (see Appendix B for code details)

## 3. Results

Across 4,812 camera trap days, we collected 49,417 images representing 20,714 terrestrial mammal encounters of 27 species. However, the number of species observed at each camera site was considerably lower than the regional species pool (mean = 7; range: 1-12). Six of these species were omitted from the analysis as they did not meet the selection criteria, or were seen only at one site (i.e. red howler monkey– *Alouatta seniculus*, pygmy squirrel*–Sciurillus pusillus pusillus,* Cotton-top tamarin*–Saguinus Oedipus,* capuchin monkey*–Cebus sp,* red-tailed squirrel*–Notosciurus granatensis,* domestic dog*–Canis lupus familiaris*). The carnivore guild was the most dominant guild, represented by eight species, followed by herbivores (*N*=6) and omnivores(*N*=4). The most common or widely distributed species, occupying more than 50% of the sites, were (in descending order): central American agouti (*Dasyprocta punctata*); common opossum (*Didelphis marsupialis*); lowland paca (*Cuniculus paca*); nine-banded armadillo (*Dasypus novemcinctus*); tamandua (*Tamandua Mexicana*); Striped hog-nosed skunk (*Conepatus semistriatus*), and; tayra (*Eira Barbara*). Conversely, less common species, occupying less than 5% of the sites, included 10 species, such as brown-eared woolly opossum (*Caluromys lanatus*), cougar (*puma concolor*), greater grison (*Galictis vittata*), among others (see Appendix A.2; C.1). Most species detected were categorized as Least Concern (LC) according to the IUCN red list (https://www.iucnredlist.org/) except for the margay (*Leopardus wiedii*), which is near threatened (NT) but was only detected in five sites out of the 89. Importantly, the critically endangered (CR) primate, tamarin (*Saguinus Oedipus*), which was not considered in the analysis, was detected at only one site, which highlights the biological importance of the area.

### Is forest integrity more important than other landscape attributes in explaining species occupancy and richness?

None of the factors examined in this study appeared to exert a substantial effect on mammal community occupancy across the Montes de Maria area, including forest cover, which goes in contrast to our predictions and suggests a wide range of differential responses within the community (Fig. 2). However, we found a moderate positive effect of forest cover on mean community occupancy (standardize beta coefficient mean = 0.361, 75% BCI = 0.053-0.669), suggesting this is probably the only relatively important driver of species habitat use in the study area. This can be further supported by the greater slope found for this variable compared to the others (mean = 0.36, sd = 0.27, Fig. 3), which were close to zero (see Appendix C.2). The effects of landscape attributes on species richness were also very weak, as indicated by the slopes or magnitude of the effects (Fig. 4).

**Figure 2.**
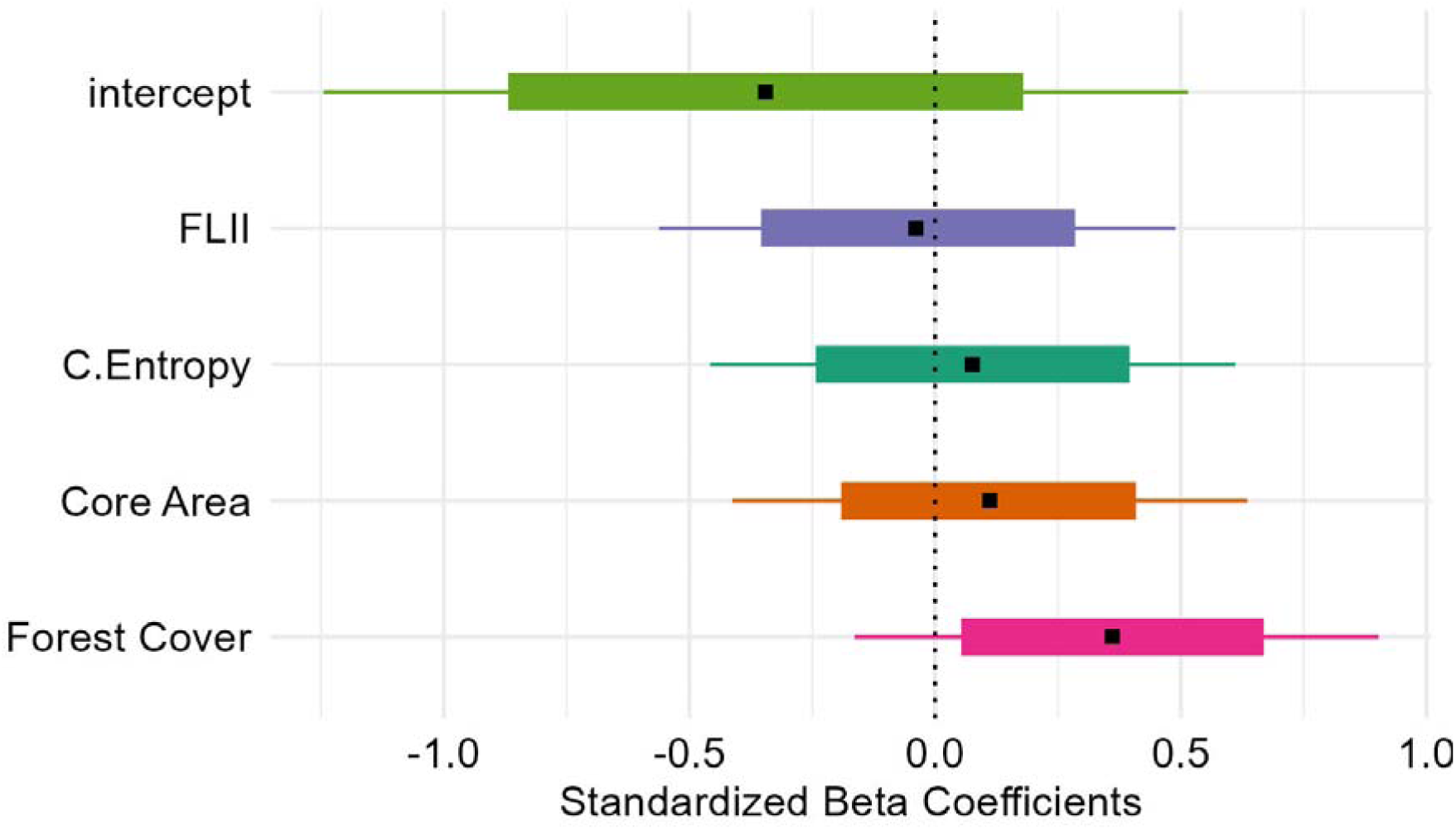
Estimated effect (slopes in logit scale) of the different variables on the mean mammal community occupancy in the Montes de Maria region, Colombia. Small black squares represent the mean, lines the 95% Bayesian Credible Intervals (BCI) and boxes the 75% BCI. Effects were considered substantial if the 95% BCI did not overlap zero and moderate if 75% BCI did not overlap zero.

**Figure 3.**
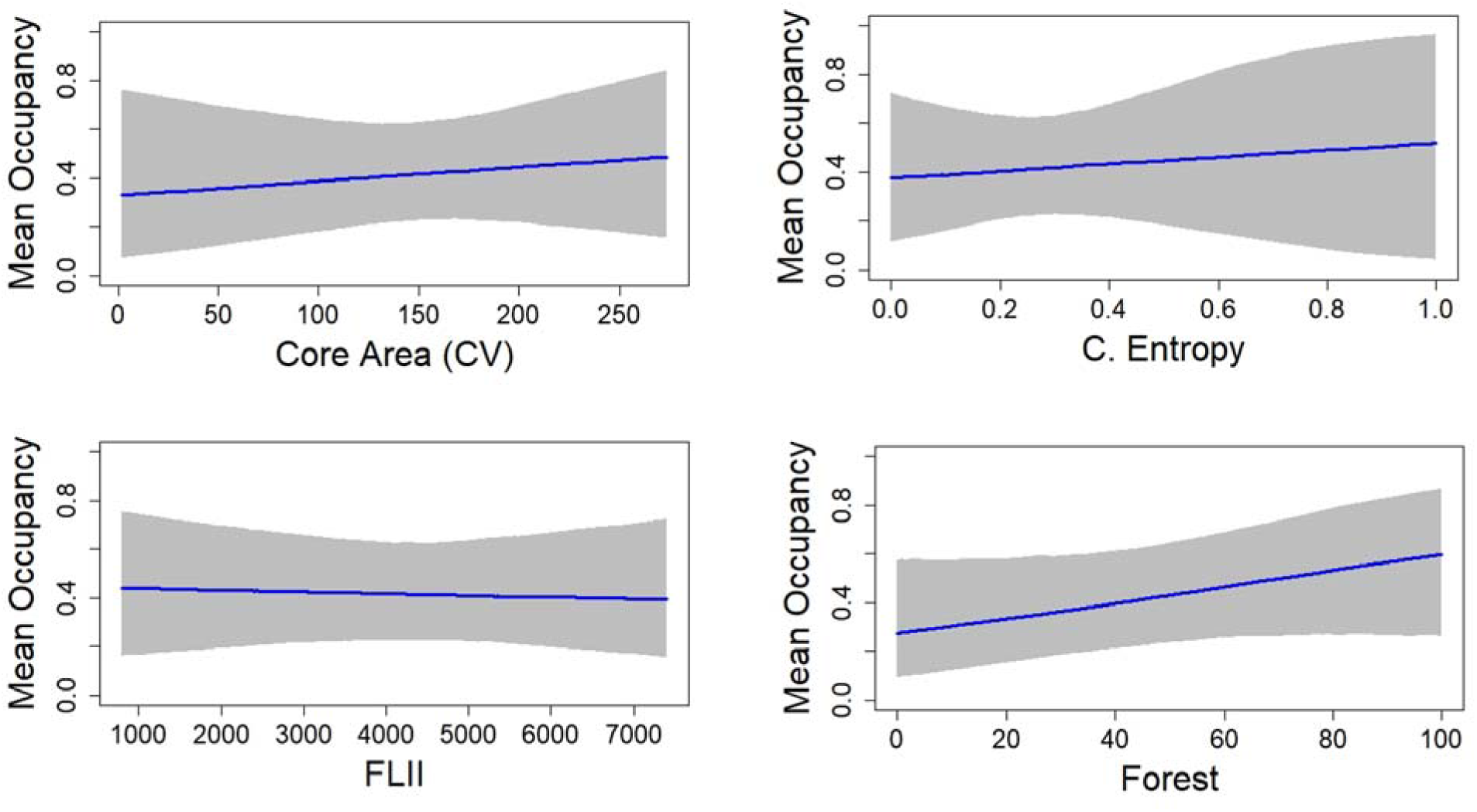
Estimated mean occupancy response of the entire mammal community in relation to four landscape variables in the Montes de Maria region, Colombia. None of the factors exerted a substantial effect but only forest cover showed a moderate effect.

**Figure 4.**
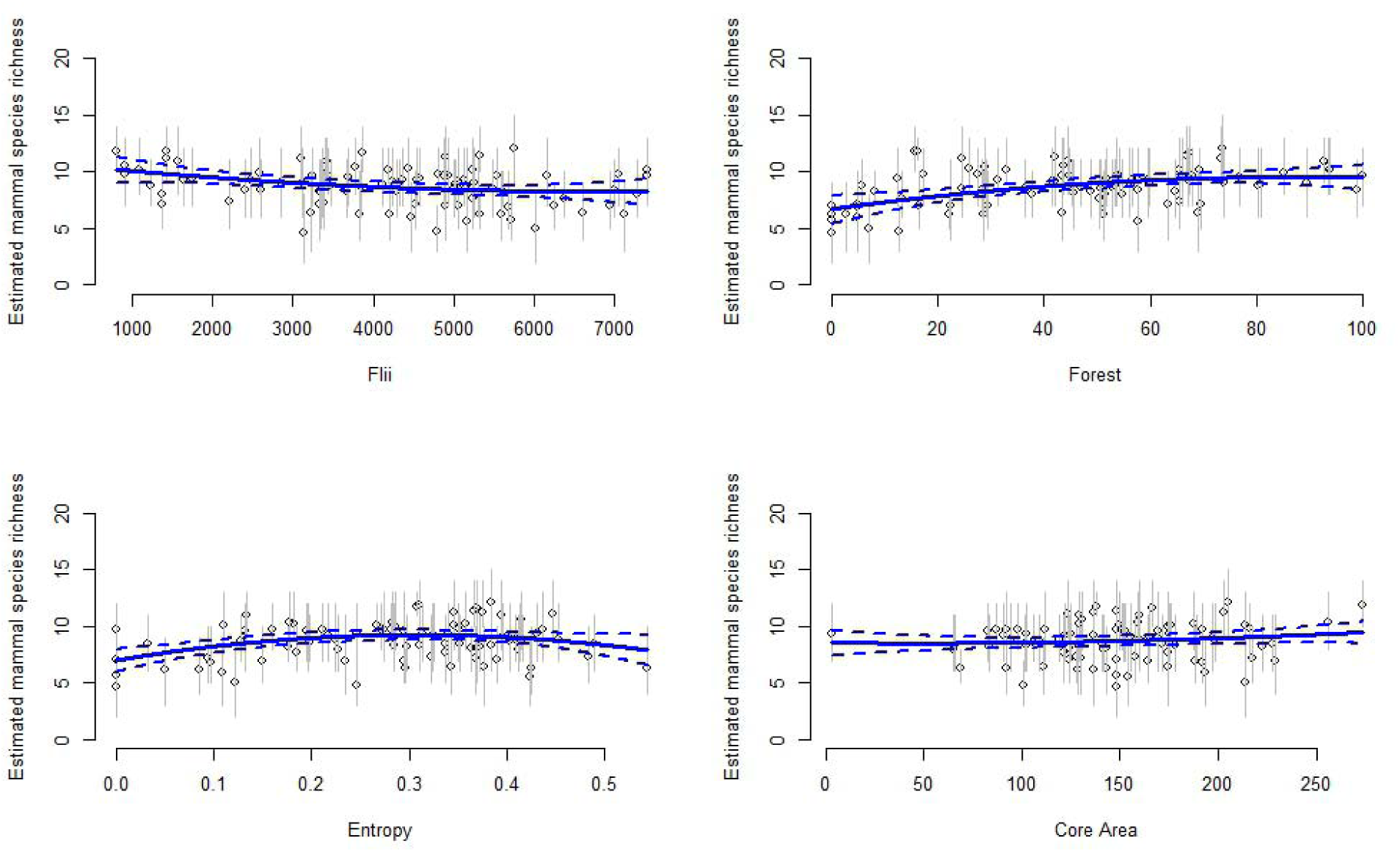
The effect of each landscape variable on estimated site-specific species richness in Montes de Maria, Colombia.

### How does the landscape influence species traits and individual species?

Similar to the results of the entire community, covariate effects within guilds were not strong in general. Forest cover was found to have a substantial positive effect on herbivores (standardized beta coefficient mean = 0.67, 97% BCI = 0.23-1.14). Similarly, the Conditional Entropy only had a moderate negative effect on omnivore species (standardized beta coefficient mean = 0.48, 75% BCI = 0.11-0.87) (Fig 5, Appendix C.2), however, its credible intervals were wide, suggesting caution when generalizing this result. This suggests, once again, that the most important driver of habitat use was forest cover and that only herbivores and carnivores are the groups where these effects are stronger in the community. When analyzing the effect of dietary breath, our results showed that forest had a moderate effect on both groups (i.e. generalist—wide diets vs specialists—narrow diets; standardized beta coefficient mean = 0.29, 75% BCI = 0.04-0.53), while no effect was found for the remaining variables (Appendix C.3). Although species with more specialist habits tended to increase habitat use as forest cover increased while generalist species had the opposite tendency (i.e. less likely to occupy landscapes with large forest cover), the magnitude of these responses was small as shown by their slopes (Fig 6). On the other hand, only forest showed a substantial effect on the occupancy of species with small home ranges (standardized beta coefficient mean = 0.35, 97% BCI = 0.009-0.69). However, the size of the effect (slope) was close to zero, suggesting there are no differences in the habitat use of species depending on their home ranges (Appendix C.5). Although the effect on species with large home ranges was slightly negative, it was no substantial, nor moderate either.

**Figure 5.**
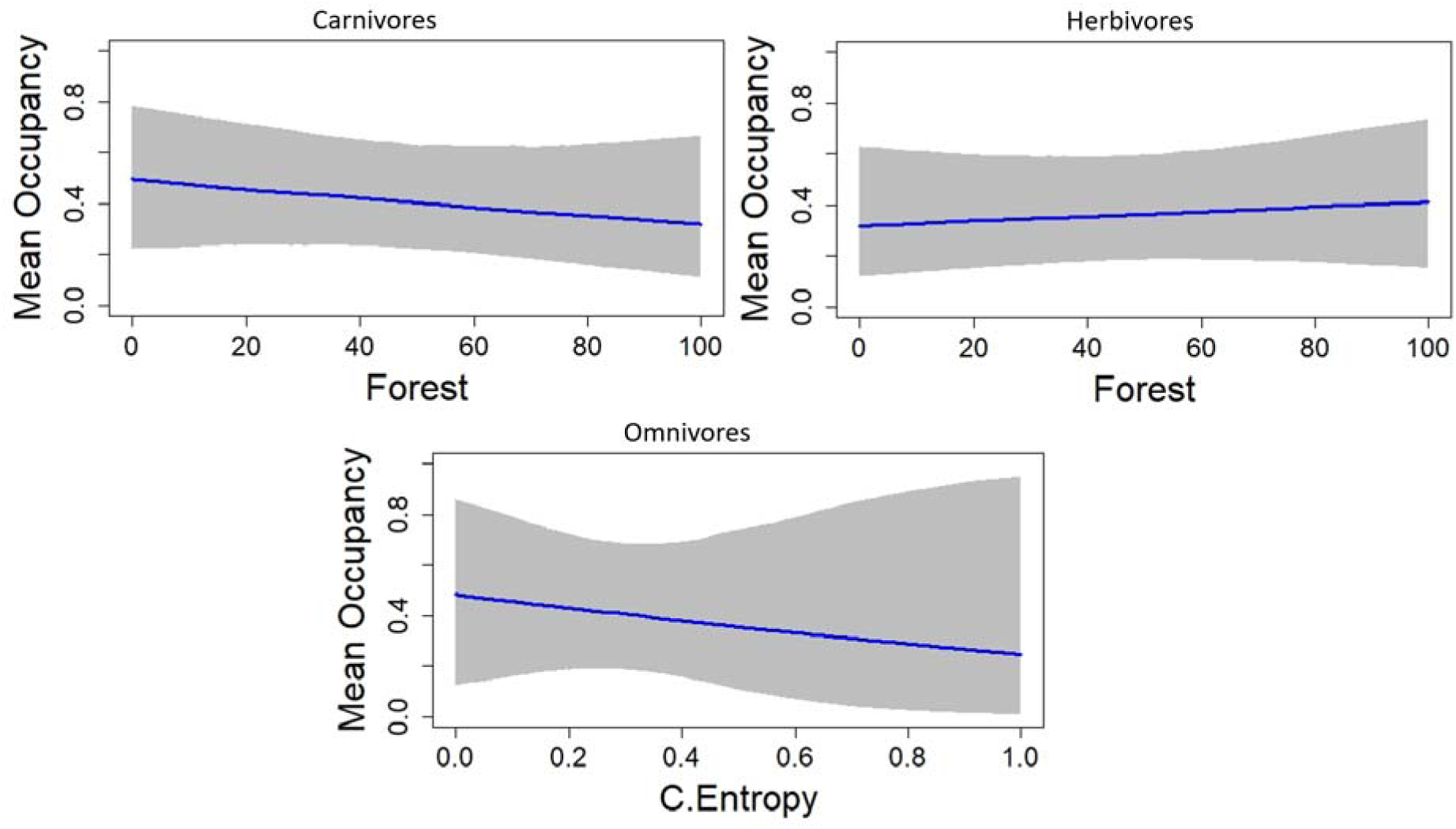
Estimated mean occupancy response of the most important variables on different guilds. Only variables and guilds with substantial and moderate responses are shown. The effect of the forest was only substantial for herbivores and not important for carnivores. The Conditional entropy only had a moderate effect on omnivore species.

**Figure 6.**
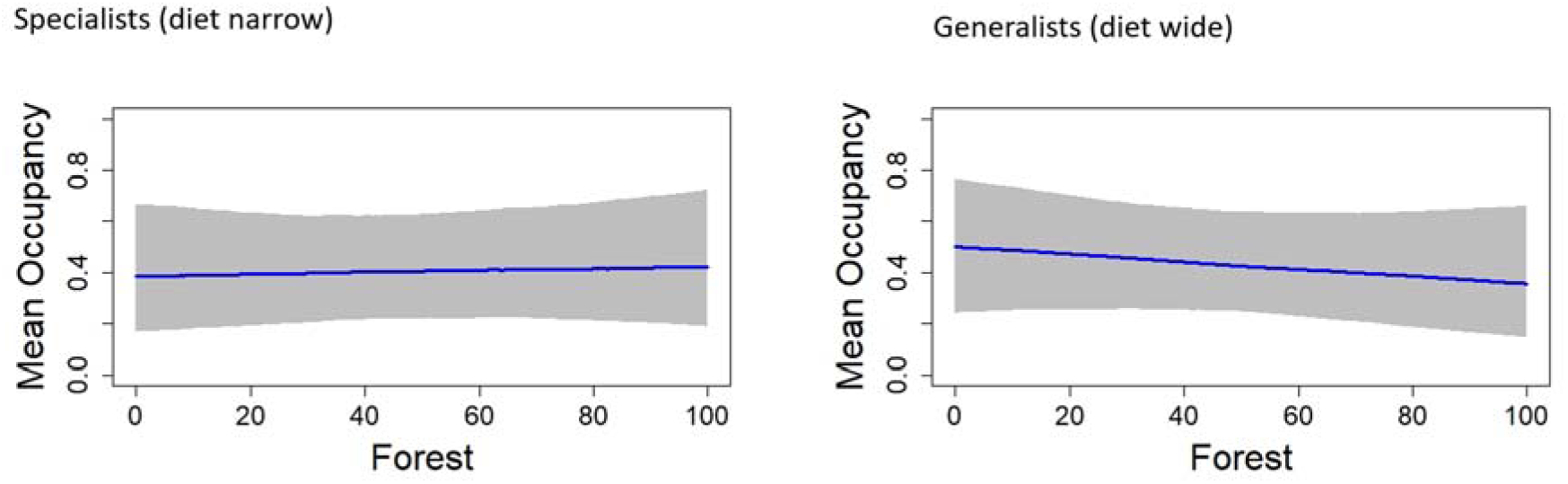
Estimated mean occupancy response of specialist (left) and generalists (right) species to forest cover in the Montes de Maria region, Colombia.

Species exhibited idiosyncratic responses to habitat covariates, though very few species showed a substantial response to some of the following variables. In general, forest cover was the only factor showing a moderate effect in several species, while the association with the other factors was only substantial or moderately important for very few species. For example, FLII had a substantial negative effect only on the lowland paca and crab-eating fox–*Cerdocyon thous*, with two species showing moderate response, crab-eating raccoon–*Procyon cancrivorus* (negative) and northern tamandua– *Tamandua mexicana* (positive). Forest was only important for the central agouti, but many species showed a moderate response (crab-eating fox, coendu–*Coendou sp*, lowland paca, tayra–*Eira barbara*, greater grison–*Galictis vittata*, ocelot–*Leopardus pardalis*, collared pecari–*Pecari tajacu*, and northern tamandua). Conditional entropy was only important for the crab-eating fox and core area for the tayra, with three species showing a moderate response (naked-tailed armadillo–*Cabassous centralis*, tayra and eastern cotton tail–*Sylvilagus floridanus*) (Fig 7, see Appendix C.6).

**Figure 7.**
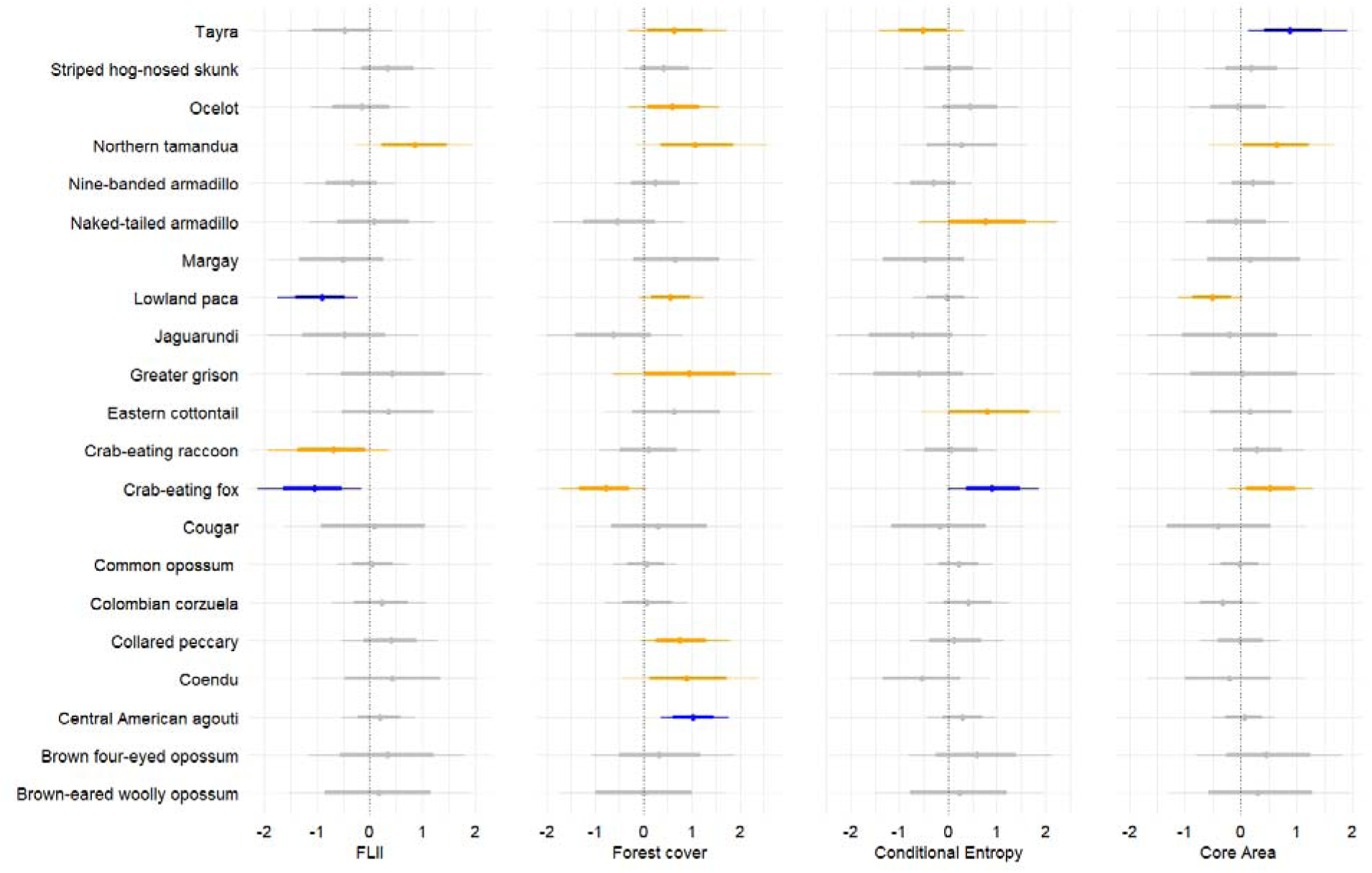
Estimated effect (slopes in logit scale) of the different variables on each species occupancy probability Montes de Maria, Colombia. The plot represents the posterior summaries (mean, 95% and 75% Bayesian Credible Intervals) of the estimated slopes on the logit-linear model for the four covariates in the occupancy (ψ) model. Points represent the mean, lines the 95% Bayesian Credible Intervals (BCI) and boxes the 75% BCI. Effects were considered substantial if the 95% BCI did not overlap zero (blue) and moderate if 75% BCI did not overlap zero (orange).

## 4. Discussion

One of the main purposes of this study was to assess the relative contribution of three prominent threats to forest-dwelling mammals (quality and extent of forest, and fragmentation processes) in determining their habitat use across an anthropogenic landscape in Colombia. Our data suggest that mammals across the Montes de Maria subregion were mostly affected by forest loss (specially the herbivore guild), but were more resistant to the degradation of forest quality and fragmentation processes (as measured by forest integrity and configuration metrics), as predicted. This is most likely due to a long history of human selective pressure or filtering process which would have resulted in only tolerant species occupying this region at current times (Newbold et al., 2018; Püttker et al., 2015). This is further supported by a lack of records of species of conservation concern, such as the jaguar (*Panthera onca*), tapir (*Tapirus terrestris*) or giant anteater (*Myrmecophaga tridactyla*), which have been reported elsewhere in the Caribbean region (Chacón-Pacheco et al., 2014; Montenegro et al., 2019). Our results confirm the capacity of some species to use landscapes with different levels of alteration and human presence, supporting other studies in anthropogenic landscapes (Boron et al., 2019; Ceballos et al., 2019; Pardo et al., 2019; Perfecto & Vandermeer, 2008). However, our results also highlight the crucial role of forest cover in the presence of mammalian species regardless of its quality, the configuration of the landscape or where patches are embedded

The influence of forest size (at patch levels or the entire landscape) compared to forest quality and other aspects, such as configuration, has been an important debate in recent years (see Arroyo-Rodríguez et al., 2020; Villard & Metzger, 2014). Fahrig, (2017), for example, suggests that the amount of forest cover in a landscape is the most important factor for maintaining species richness regardless of the fragmentation process (e.g. geometry, number, area, and location of individual patches; see “habitat amount hypothesis”). Our results align with this hypothesis but in relation to habitat use of a mammalian community since our configuration proxy (e.g. core area, conditional entropy) did not show a substantial nor moderate effect. Interestingly, though, our results showed that the effect of the forest cover was only moderate, except for herbivores. The importance of forest cover is reflected in other studies that have found the existence of tipping points or thresholds in reducing forest cover that once surpassed can trigger drastic declines in populations (e.g. Roque et al., 2018).

A potential reason as to why there was only a moderate response to forest cover might be related to the quality of the matrix. Although we did not measure the quality of the matrix per se, studies suggest that biodiversity extinction thresholds also depend on matrix type (Boesing et al., 2018). According to this hypothesis, if the quality of the matrix is good, then the need for more forest cover is less important for maintaining species. Therefore, it is possible that the studied landscape does not have strong connectivity issues, and the matrix might be relatively permeable, allowing animals to move across the sites or patches. Furthermore, different matrix types may offer different levels of resources, which, generalist species, such as those left in the community, may be able to exploit, uncoupling their dependence on forest resources. The study area is a mosaic of many different agricultural products such as cocoa, ñame, yuca, and different fruits and vegetables, most of them cultivated at small scales. Evidence suggests that some species are not strongly limited by some of these crops as animals can use them to move across the landscapes, particularly generalist carnivores and deer species (e.g. López-Ramírez et al., 2020; Nogeire et al., 2013). As a matter of fact, 62 interviews conducted in the area showed that people have conflicts with some mammal species such as the tayra, hog-nosed skunks (*conepatus semistratus*), jaguarundi and others (IAvH, 2018), which suggest that these species use anthropogenic mosaics. A further study is required to quantify how mammal species are using the anthropogenic mosaics in this region.

### Functional groups

Evidence suggests that anthropogenic landscapes tend to, not just increase taxonomic homogenization, but could also drive genetic and functional homogenization (Olden et al., 2004). In this study, functional groups (dietary breath, home range, or trophic group) were generally resilient to changes in habitat extent, quality or configuration, except for the herbivore guild. In fact, the assemblage was well represented by different functional groups, except for large-body mammals. Our results contradict Regolin et al. (2021), who found that other proxies of habitat quality (distance to waterbodies and abundance of high shrubs) were more important than habitat amount in determining herbivores and frugivorous mammals’ habitat use in the Brazilian Pantanal. It is worth noting that in their study, they also used some proxies for habitat quality at finer scales, such as the proportion of food resources (i.e. plants), while in our study, the proxy for habitat quality is a landscape measure that represents structural integrity. As such, although FLII represents the degree to which forest structure, composition and function have been modified by anthropogenic actions, it uses other proxies not related to microhabitat or specific structural indices of the forest (e.g. observed human pressures and inferred human pressures).

Other studies have found that species with specialized niches are usually more vulnerable to habitat loss, fragmentation, and other disturbances (Purvis, 2000). In our study, species with wide or narrow diets (generalists and specialists, respectively) did not respond strongly to any of the variables and only showed a moderate response to forest loss, suggesting that species across this landscape are not filtered by their ecological niche. As expected though, generalist species tended to occupy areas with less forest cover. In some regions across the tropics, insectivores and frugivores mammals exhibit high sensitivity to land-use change and other human disturbances (Rovero et al., 2019). However, in our study area, the effect of the remaining forest was only strong and positive for herbivores. Our results also go in accordance with the findings of Bedoya-Durán et al. (2021) in the Andean region of Colombia. They examined the influence of some landscape attributes (i.e. land use type, fragmentation, connectivity, and human disturbance) on the occupancy of species within functional groups (i.e. guilds, size, and niche breadth) across privately protected areas and non-protected sites and found no clear effect, but suggest that the occupancy of forest-restricted species and large species tended to be positively related to the proportion of forest at a site. Boron et al., (2019) also found weak relationships with several variables in a community in the Magdalena Medio in Colombia, except for forest cover that strongly modulated habitat use of species. We did not include species size as a covariate as we only detected one large species (puma–*Puma concolor*) and two small species (Brown-eared woolly opossum–*Caluromys lanatus*; Brown four-eyed opossum– *Metachirus nudicaudatus*), which were also very rare in the landscape and could, therefore, bias results. However, we can speculate that the lack of substantial effects of the variables on each of the examined guilds, and the lack of detection of large species (tapir, jaguar, giant anteater) suggest that a possible potential filtering in this anthropogenic landscape is mostly driven by size, whereby large species are disappearing as suggested by Suraci et al. (2021).

The ecological plasticity of carnivore species in anthropogenic landscapes has been documented, particularly mesocarnivores such as foxes and jaguarundis in industrialized agricultural lands (Boron et al., 2019; Pardo et al., 2018b), other heterogeneous landscapes (Daily et al., 2003) and even cities (Hansen et al., 2020). In our study, forest cover showed no effect on carnivores, which suggests that carnivores are more tolerant to other land uses and occur in the landscape regardless of their characteristics. This further suggests that there are other more important drivers of carnivore occupancy. For example, carnivores might select habitats based mostly on prey availability and/or catchability, rather than habitat characteristics. Although forest cover was not a driver of insectivores’ habitat use, the direction of the effect was positive, highlighting the importance of forests for the conservation of this group.

Although the effect of forest cover was only moderately important in species with small home ranges and not relevant for species with large home ranges, the slopes suggest that species with small dispersion movements are distributed almost evenly in the landscape regardless of the size of the forest. In contrast, species with large home ranges tended to occur more in areas with little forest cover with decreasing habitat use when forest cover increased. This was an unexpected finding as species with large home ranges are expected to move across the landscape more easily and regularly. All species with large home ranges were carnivores (tayra, ocelot, margay, jaguarundi, cougar), therefore this negative relationship with forest cover might suggest that good quality resources or prey are not restricted to large forest patches, and that small patches might be providing good resources (including human-made food such as crops, domestic species). On the other hand, there could also be a detectability issue, since carnivores are species with large home ranges, the chances of detecting these species in large forest patches could be lower. This is because there would be more space for them to move freely reducing the likelihood of crossing in front of the cameras, while in small patches the options are more restricted and therefore the chances of being detected by the camera when the animal is nearby are greater. This result on carnivores might suggest that the carnivore species are most likely moving across small patches, either because there are no connectivity issues or because of their capacity to move across different habitat types, including disturbed habitats which could provide food resources (see above paragraph).

### Species-specific responses

When trying to understand which species are mostly driving the occupancy patterns of the community, we found only a few species substantially affected by the landscape covariates, but some contradicted our predictions. For example, we expected that the quality of the forest would substantially drive the occupancy of lowland paca since this is a forest specialist species. Our results confirmed this, but unexpectedly this relationship was found to be negative, while forest cover showed a moderate positive association. Since Core Area also had a strong effect, it is apparent that the lowland paca distribution is not affected by edge effect issues, as opposed to the tayra that was strongly and positively affected by this variable and tended to increase habitat use in more homogeneous landscapes. Although tayra is a relatively common species in anthropogenic landscapes, our results highlight that they tend to occupy larger interior habitats inside the forest, which goes in accordance with Bianchi et al. (2021) findings in a heterogeneous landscape in Brazil. The crab-eating fox habitat use was greater in areas with low forest integrity and more heterogeneous composition, confirming its flexibility to move across anthropogenic landscapes (Boron et al., 2019; Pardo et al., 2019). The Central American agouti was clearly positively influenced by forest cover despite its quality which also goes in accordance with other studies, highlighting their dependency on habitat cover (Boron et al., 2019; Pardo et al., 2018). The presence of these forest-restricted species (including those not used in the analysis, see methods) suggests that some vital process such as seed dispersal is still occurring in the area helping to maintain the forest dynamic (see Camargo-Sanabria et al., 2014). However, the absence of large mammals limits the contribution of the present fauna as this group of species has important roles as, for example, the regulatory role of apex predators, and the dispersal of large-seeded plants which limits the growth of large wooded trees and in consequence limits carbon storage capacity of the forest (Bello et al., 2015).

### Conservation

Our study emphasizes the resilience of several tropical mammals in human-modified landscapes but highlights the importance of conservation interventions that preserve or restore forest extent in human-modified landscapes. Restoration of forests and other ecosystems has become a key strategy for the United Nations (UN). As a matter of fact, the UN has declared that we are currently in the decade of restoration (2021-2030, see https://www.decadeonrestoration.org/) where considerable emphasis is being placed on restoring forests. Although restoration projects include many aspects (e.g. expanding forest, restore ecosystem services, improving peoplés livelihoods, etc.), due to the alarming rates of deforestation expanding forests (particularly in formed forested land) has become a major interest. Our results support the importance of this strategy and suggest that priority should be given to maintaining current patches of forest and increasing its availability for mammals. As suggested by the UN, abandoned or non-productive land could be an important opportunity to fulfil this purpose. Some scholars suggest that to optimize biodiversity conservation in human-modified landscapes, forest patches should account for at least 40% of the landscape and these patches should be embedded in a high-quality matrix or with less agriculture (Arroyo-Rodríguez et al., 2020). Our results support the claim to protect as much forest as possible but suggest that extant species are not restricted by forest quality. Therefore, we believe that conservation strategies in the study area should focus on conserving as much forest as possible (e.g. avoiding further deforestation) regardless of its conservation status or quality, configuration of landscape or size/shape of individual patches; at least for non-threatened species, such as those currently found in these transformed landscapes.

The strong association of herbivores with forest cover suggests their high vulnerability to the reduction of forest cover, which suggests further attention. This is supported by other studies that highlight the importance of maintaining forest cover along gradients of human land use, given the evidence for stronger negative effects of forest loss on biodiversity in comparison to other factors such as forest configuration or other fragmentation processes per se (see Fahrig, 2017). In this sense, private conservation could play an important role in anthropogenic landscapes, as evidence suggests that secondary forests or even small patches have important conservation implications (e.g. Fahrig, 2017). Even more, the presence of species of conservation concern such as the margay and the tamarin, confirms the importance of Montes de Maria landscape for the conservation of species. The loss of herbivores can have important implications for biodiversity integrity as this group is not just important for maintaining forest conditions, but also because of their role as prey for larger carnivores.

On the other hand, as suggested by Hulme-Beaman et al. (2016), occupation does not denote dependency or that the species are not facing any threats. It is also possible that apparently stable populations are at their maximum tolerance, and some could be experiencing a delayed response to threats (see Semper-pascual et al., 2017). Only a multitemporal analysis could provide insights into the potential conservation risks for the species currently occupying the study area. While some species might have shown tolerance to this landscape, this apparent tolerance might be limited to a certain level of alteration as there are thresholds for these responses, that once passed, the direction and magnitude of these effects could be dangerous (Pardo, et al., 2018; Suraci et al., 2021). Future studies should explore the possible impact of hunting and domestic animals in driving species distribution in this landscape, which could even be stronger than landscape measures (e.g. Deere et al., 2020). The lack of detection of species of conservation concern (jaguar, tapir, and giant anteater) suggests that they might have undergone local extinction or simply that they are so rare that they went under-detected. However, the area still supports potential prey for large predators (collared peccary, deer, lowland paca, etc.). Since we detected puma (another large predator), it could be speculated that the absence of jaguars could be mainly caused by poaching, which is considered one of the main factors threatening this species.

In conclusion, our results reinforce the need to conserve any type of forested areas for the conservation of mammals regardless of the extent of human-driven habitat modification (integrity) or the current configuration of the surrounding landscapes (fragmentation process). Our multispecies occupancy model allowed us to investigate the entire community, functional groups and species (some of them hard to study on their own due to low detectability) and allowed us to see the idiosyncratic responses of mammals to fragmentation processes. We suggest that variation across species must be taken into account when attempting to make inferences for an entire assemblage or functional groups, as the effect can be contrasting even within these groups.

## 5. Acknowledgements

## Notes

### Competing Interest Statement

The authors have declared no competing interest.

### Summary of Updates

figure 1 was updated few lines in the introduction and discussion were reorganized

